# Revisiting Endothelial Tropism of SARS-CoV-2 Using a Cell-Specific hACE2 Mouse Model

**DOI:** 10.1101/2025.05.01.651662

**Authors:** Sahine Lameire, Nincy Debeuf, Julie Deckers, Caroline De Wolf, Manon Vanheerswynghels, Wendy Toussaint, Lize De Vlieger, Lien Van Hoecke, Arnout Bruggeman, Sieglinde De Cae, Bert Schepens, Stijn Vanhee, Bart N. Lambrecht

## Abstract

Severe COVID-19 is frequently associated with vascular complications, raising ongoing debate about whether SARS-CoV-2 can directly infect endothelial cells and thereby contribute to disease pathogenesis. Although endothelial cells express angiotensin-converting enzyme 2 (ACE2), the *in vivo* relevance of endothelial-restricted viral tropism remains unclear. To directly assess the consequences of endothelial-restricted SARS-CoV-2 tropism *in vivo*, we generated a transgenic mouse model expressing human ACE2 under control of the endothelial-specific *Cdh5* promoter (*Cdh5*-hACE2). Despite confirmed pulmonary endothelial expression and protein presence of hACE2, SARS-CoV-2 infection of *Cdh5*-hACE2 mice did not induce clinical illness, detectable viral replication, immune cell influx in the lung, or histopathological abnormalities in the lung or brain. These findings indicate that endothelial-restricted SARS-CoV-2 tropism alone is insufficient to drive productive infection and clinical disease *in vivo*, suggesting that endothelial involvement in COVID-19 likely arises in the context of broader cellular infection or systemic host responses rather than from primary endothelial infection.

## Introduction

Severe acute respiratory syndrome coronavirus 2 (SARS-CoV-2) is the causative agent of COVID-19, which escalated into a global pandemic in 2020. SARS-CoV-2 is an enveloped, positive-sense, single-stranded RNA virus belonging to the genus *Betacoronaviridae* (1). The severity of COVID-19 cases varies from asymptomatic to those experiencing severe acute respiratory distress syndrome (ARDS) with possible fatal outcomes, as well as post-acute sequelae including long COVID (2, 3).

Whereas SARS-CoV-2 initially replicates in the upper airways, the clinical manifestations associated with severe COVID-19 predominantly involve the lower respiratory tract, including pneumonia, cough, dyspnea, and ARDS (4). Early during the pandemic, however, COVID-19 rapidly emerged as a multi-organ disease, as patients presented with a broad spectrum of extrapulmonary symptoms such as diarrhea, proteinuria, neurological complications, thromboembolic events and cardiovascular inflammation (4).

A growing body of evidence has implicated vascular pathology as a unifying feature underlying these diverse clinical manifestations. Among the hallmarks of severe COVID-19, endothelial cell injury and dysfunction, collectively referred to as endotheliopathy, have consistently been associated with disease severity and mortality (5). Initial insights into endothelial involvement stemmed largely from autopsy studies, which revealed widespread microangiopathy and thrombosis in the pulmonary vasculature of COVID-19 patients, occurring at markedly higher frequencies than in influenza-associated lung disease (6). Endothelial damage in the lungs was further linked to perivascular immune cell infiltration, suggesting a close interplay between viral infection, inflammation, and vascular injury.

Importantly, endothelial dysfunction is not restricted to the pulmonary circulation alone. Microvascular injury and thrombosis have been reported in multiple distal organs, including the kidneys, heart, and brain, and are thought to contribute to organ-specific complications such as proteinuria, renal failure, cardiovascular dysfunction, and neurological sequelae (7, 8). Together, these findings support the concept that SARS-CoV-2–induced vascular pathology plays a central role in multi-organ dysfunction and the adverse clinical outcomes.

Despite the strong association between vascular pathology and severe COVID-19, the mechanisms driving SARS-CoV-2–induced endothelial dysfunction remain controversial. Initial studies largely attributed endothelial injury to indirect effects of infection, whereby excessive inflammation originating from infected epithelial and innate immune cells generates a systemic pro-inflammatory milieu that secondarily disrupts endothelial integrity (9–11).

Other evidence emerged suggesting that SARS-CoV-2 may also directly interact with endothelial cells (12). Viral particles or the spike (S) protein alone have been reported to alter endothelial function, promoting inflammation and microvascular dysfunction even in the absence of productive infection (13, 14). However, conflicting data persist regarding true endothelial tropism and the ability of SARS-CoV-2 to replicate within endothelial cells. As a result, the relative contribution of indirect inflammatory injury, spike-mediated endothelial activation, and bona fide endothelial infection to COVID-19-associated vascular pathology remains unresolved.

To specifically interrogate the contribution of endothelial-restricted SARS-CoV-2 interactions to clinical disease, we generated *Cdh5*-hACE2 mice, in which hACE2 expression is driven by the endothelial *Cadherin-5* promoter. This model was designed to isolate endothelial-restricted SARS-CoV-2 entry in the absence of epithelial infection. Using this endothelial-restricted hACE2 model alongside the widely used *Krt18*-hACE2 mice, we performed an in-depth comparative pathological analysis following SARS-CoV-2 infection, assessing clinical disease, viral replication, immune cell influx, and histopathological changes across multiple organs.

## Results

### Endothelial-specific hACE2 expression in *Cdh5*-hACE2 mice supports Spike protein binding

To investigate the debated role of endothelial cells in SARS-CoV-2 infection, we generated a transgenic mouse model with endothelial-restricted hACE2 expression driven by the *Cdh5* promoter. A detailed description of the generation of the transgenic line is provided in the Materials and Methods section. This model was compared with the widely used *Krt18*-hACE2 mouse, the current reference model for SARS-CoV-2 infection in mice (Fig.1A,B). Both quantitative PCR (RT-qPCR) and Western blotting showed clear and significant expression of hACE2 in the lungs of all transgenic mice in comparison to non-transgenic wild type littermates (WT) (Fig.1C,D). RT-qPCR of fluorescence-activated cell sorted (FACS) lung cells confirms the expected endothelial cell-specific expression in the *Cdh5*-hACE2 tg mice (Fig.1E). While in *Krt18*-hACE2 mice, hACE2 expression is driven predominantly in epithelial compartments, we pick up significant expression in endothelial cells as well. However, flow cytometric staining of the hACE2 receptor on endothelial cells with a fluorescently labelled Spike-protein only shows significant staining in the *Cdh5*-hACE2 tg mice.

**Fig 1:**
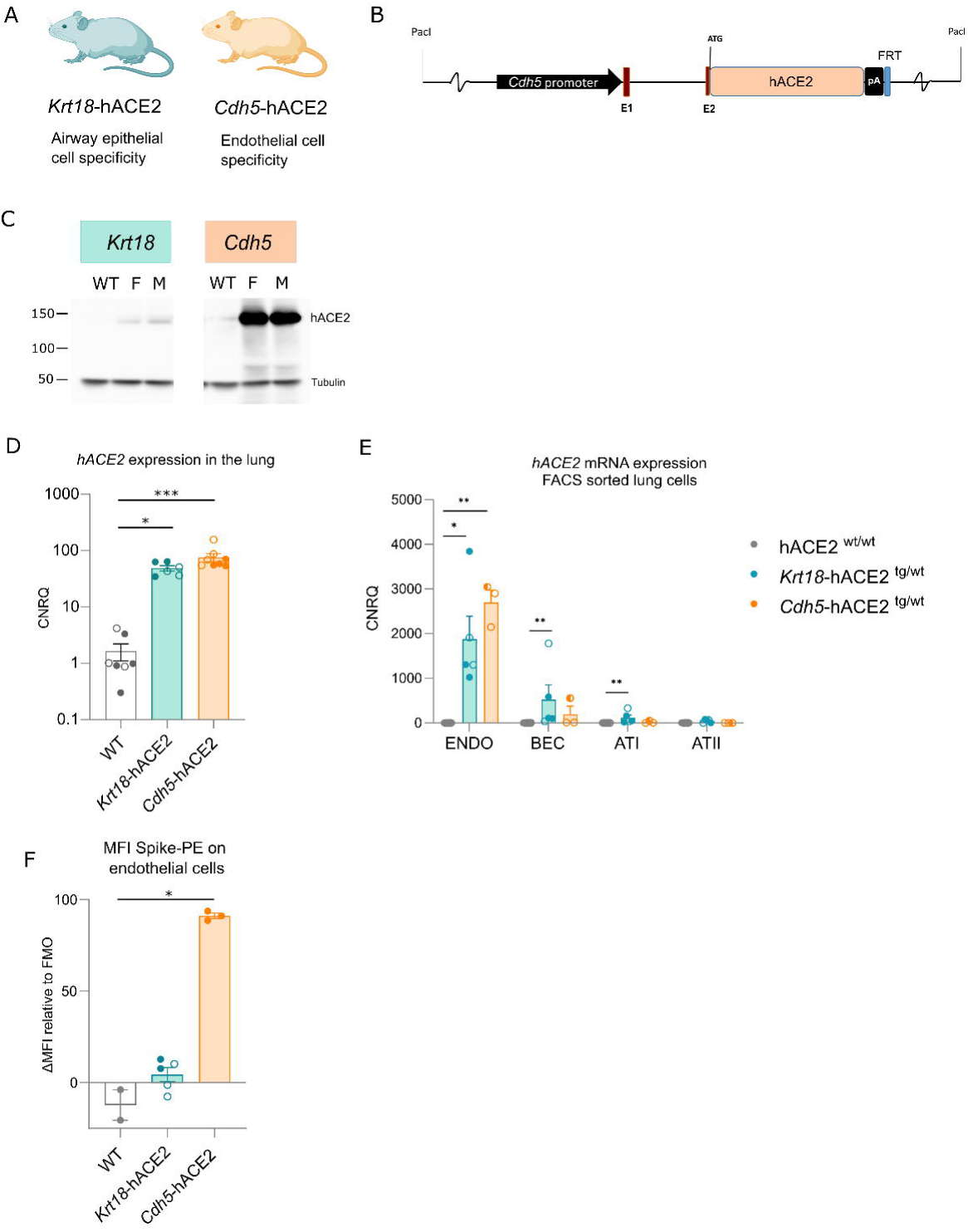
**A.** Schematic overview of the hACE2-transgenic mice with their corresponding primary expected tissue expression of the hACE2 receptor. **B**. Graphical representation of the constructs used to generate the *Cdh5*-hACE2 transgenic line. **C.** Western blot analysis of hACE2 protein in the lung of steady state hACE2-transgenic mice and accompanying β-tubulin loading controls. Samples were normalized to the same protein concentrations before loading on gel. WT = wildtype, F = female, M = male. **D.** RT-qPCR analysis of *hACE2* mRNA expression in lung tissue of steady-state hACE2 transgenic mice. **E.** RT-qPCR analysis of *hACE2* mRNA expression in FACS sorted lung cells of steady-state hACE2 transgenic mice. ENDO: endothelial cells, BEC: Bronchial-epithelial cells, ATI: alveolar epithelial type 1 cells, ATII: alveolar epithelial type 2 cells. **F.** PE-Spike protein binding on endothelial cells is shown as ΔMFI relative to the corresponding fluorescence-minus-one (FMO) control. Statistics: Data are shown as means ± SEM. Male mice are displayed in filled dots while females are open dots, pooled samples of both genders are represented as half-filled dots. After assessing normality by a Shapiro-Wilk normality test, non-parametric data were analysed with a Kruskal-Wallis test, where each transgenic line was compared to WT. ∗p < 0.05; ∗∗p < 0.01; ∗∗∗p < 0.001.

### Absence of clinical disease, viral replication, and lung pathology in SARS-CoV-2–infected *Cdh5*-hACE2 mice

We first performed titration experiments to find the optimal inoculum dose of the European D614G strain (EVAg) in *Krt18*-hACE2 mice, a fully established mouse model for COVID-19 (15–18). Intratracheal (i.t.) infection with 450 plaque forming units (pfu) resulted in *Krt18*-hACE2 mice displaying an average reduction of 8,06% ± 1.65 in their body weight at 7 dpi accompanied by respiratory distress and reduced mobility (Fig.2A,C). By 11 dpi, when the survivor mice had regained their initial weight, only 50% of *Krt18*-hACE2 mice had survived (Fig.2A). By contrast, *Cdh5*-hACE2 mice, with hACE2 expression restricted to endothelial cells, did not lose weight nor displayed clinical signs of disease when infected via inhalation of the same dose (450 pfu) of viral inoculum, a phenotype that persisted even after a 100-fold increase in viral dose (Fig.2A,C). Non-transgenic wild type littermates were infected with an identical viral dose as control, and were not susceptible to infection. To circumvent the fact that lung tissue damage might be necessary for the virus to reach the hACE2 viral entry receptor on the endothelium, we bypassed this barrier by also infecting a separate group of *Cdh5*-hACE2 mice intravenously with a very high viral dose (10^4^ pfu). However, intravenous infection still did not cause clinical illness in these mice (Fig.2B).

**Fig 2:**
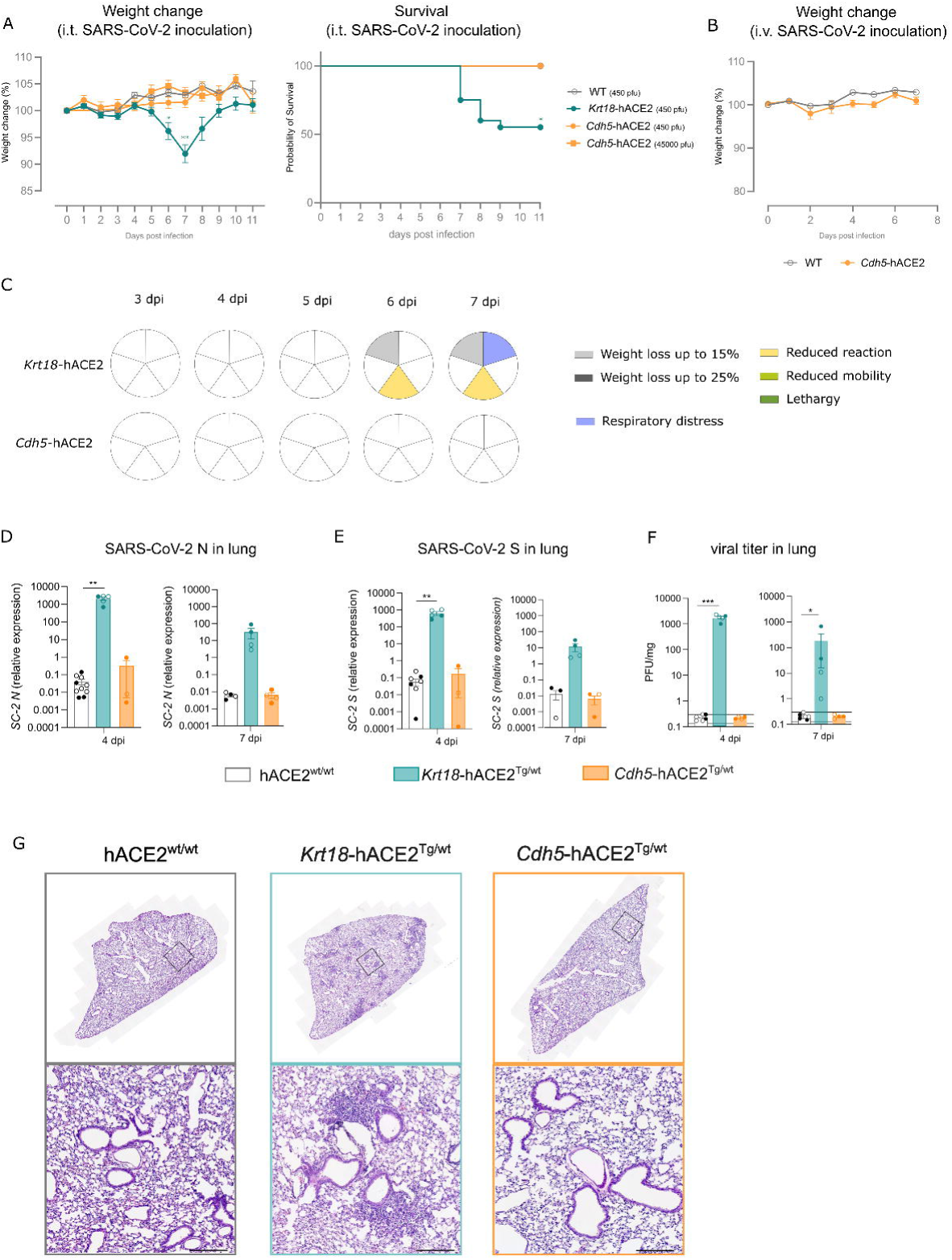
**A.** Weight loss and survival curve of hACE2-transgenic mice (6-12 weeks old) after intratracheal SARS-CoV-2 infection (450 pfu). Data represent 6-20 mice per group, pooled from three separate experiments, and are shown as means ± SEM. **B**. Weight loss of *Cdh5*-hACE2 mice (6-12 weeks old) after intravenous SARS-CoV-2 infection (10^4 pfu). Data represents 6 mice per group and are shown as means ± SEM. **C**. Schematic overview of the clinical signs observed post intratracheal SARS-CoV-2 infection with 450 pfu. Each row represents a hACE2-transgenic line and columns represent the number of days post infection. Weight-loss and lethargic symptoms are gradual measurements, where a darker shade represents worse symptoms. Respiratory distress is a discontinuous measurements meaning if the mice experienced the signs, the pie is filled. The pie represents both males and females. **D/E**. RT-qPCR results of N- and S-gene expression in post-caval lung lobe of SARS-CoV-2 infected hACE2 transgenic mice (i.t., 450 pfu) 4 and 7 days post infection. The virion associated N gene represents total viral RNA present in the sample (relative to an internal standard), regardless of whether the virus is infectious or not, while the S-gene represents viral gene expression relative to house-keeping-genes. Male mice are displayed in filled dots while females are represented with open dots. **F:** Determination of infectious viral progeny by plaque assay on VeroE6/TMPRSS2 cells of homogenised lung samples of SARS-CoV-2 infected hACE2 transgenic mice (i.t., 450 pfu) 4 and 7 days post infection (PFU/mg). PFU, plaque-forming units. Male mice are displayed in filled dots while females are represented with open dots. Black lines indicate the sample-specific plaque assay limit of detection converted to PFU/mg as LOD95%=12/M (M in mg; assay LOD = 24 PFU/mL; homogenization volume = 0.5 mL). Nondetects are plotted at their own LOD. **G**. Representative H&E-stained lung sections on 7 dpi. Scale bar = 200 µm. Statistics: After assessing normality by a Shapiro-Wilk normality test, parametric data were analysed with a One-way ANOVA and non-parametric data were analysed with a Kruskal-Wallis test. The statistics of the survival data was tested with the Kaplan-Meier test. Each transgenic line was compared to WT. ∗p < 0.05; ∗∗p < 0.01; ∗∗∗p < 0.001.

Despite the absence of overt clinical disease, we next quantified viral RNA at the tissue level using qPCR. We collected tissues at day 4 post-infection to assess early viral dynamics before overt disease, and at day 7 post-infection, corresponding to the stage at which *Krt18*-hACE2 mice develop more severe weight loss and begin to succumb to infection.

Expression of the virion-associated N gene was only detected in the lungs of *Krt18*-hACE2 mice (Fig.2D). Because this signal may reflect residual input virus, we additionally quantified S-gene RNA to further assess viral presence in the tissue (Fig.2E). Again, S-gene was highly detected in *Krt18*-hACE2 mice but no signal was picked up in the lungs of the *Cdh5*-hACE2 mice. To distinguish and quantify infectious viral particles (viral titer), a plaque assay on VeroE6/TMPRSS2 cells was performed (Fig.2F). Samples were weighed post-collection for normalisation. Assay sensitivity was defined by the Poisson 95% limit of detection (LOD□□), calculated from the total undiluted-equivalent volume plated. In *Krt18*-hACE2 mice, the viral titer in the lung was strongly elevated at 4 dpi and was drastically reduced by 7 dpi.

On histological examination of the lungs one day prior to reaching the anticipated humane endpoint, we observed perivascular and peribronchial immune cell infiltration exclusively in the lungs of the *Krt18*-hACE2 Tg mice (Fig.2G). Despite clear expression of hACE2 in lung endothelium, Cdh5-hACE2 mice were not susceptible to infection, nor did they develop vascular problems.

### Immune cell influx in the lung upon SARS-CoV-2 infection is dependent on the hACE2 expression pattern

Although no viral replication or overt lung pathology was detected in Cdh5-hACE2 mice, we next assessed whether subtle pulmonary immune alterations were present. Therefore, we performed multicolour flow cytometry on lung tissue and quantified the number of infiltrating immune cells, while adhering to the stringent biosafety measures imposed in a Biosafety Level 3 lab. At 7 days post infection (dpi), *Krt18*-hACE2 transgenic female mice exhibited a marked increase in pulmonary neutrophils. In parallel, these mice showed a pronounced influx of CD88□CD11b□Ly6C□ monocytes into the lung, accompanied by a reduction in SiglecF□CD11c^hi^ alveolar macrophages (AMs) (Fig. 3B-C). This AM “disappearance” reaction after viral infection has been described before (19–21). At 7 dpi, CD8 T cells, and not CD4 T cells, were significantly elevated in *Krt18*-hACE2 mice compared to wildtype mice (Fig.3D-E). This was accompanied by robust IFN-*γ* production in the lung, suggesting activation of the infiltrating T cells (Fig.3F). Immune cell numbers were not altered in lungs of *Cdh5*-hACE2 mice compared to wildtype mice (Fig.3A-F).

**Fig 3:**
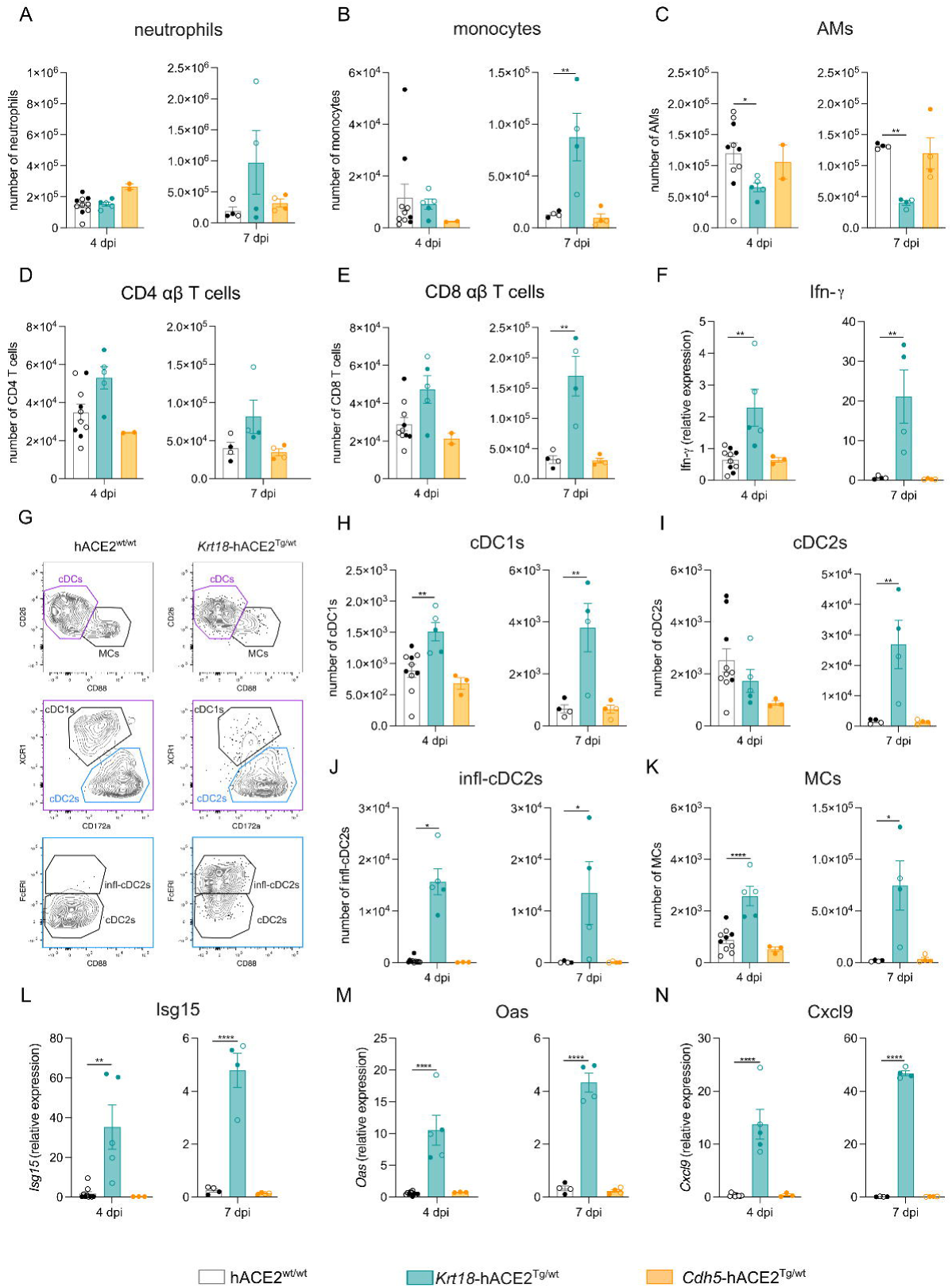
**A-E, H-K.** Flow cytometric measurements of inflammatory cells in the lung of SARS-CoV-2 infected mice (i.t., 450 pfu) 4 and 7 days post infection. Data are shown as means ± SEM. Male mice are displayed in filled dots while females are represented with open dots. **G.** Gating strategy to distinguish DC subsets presented in H-K **F, L-N**. Relative mRNA expression of *Ifn-γ*, *Isg15*, *Oas* and *Cxcl9* in the post-caval lung lobe of SARS-CoV-2 infected hACE2 Tg mice (i.t., 450 pfu) 4 and 7 days post infection. Statistics: After assessing normality by a Shapiro-Wilk normality test, parametric data were analysed with a One-way ANOVA; and non-parametric data were analysed with a Kruskal-Wallis test. Each transgenic line was compared to WT. ∗p < 0.05; ∗∗p < 0.01; ∗∗∗p < 0.001; ∗∗∗∗p < 0.0001.

Next, we zoomed in on dendritic cells (DCs), as these are the cells responsible for CD8 and CD4 T cell priming. A gating strategy was employed to distinguish monocyte-derived cells (MCs) (CD88^+^) from conventional DCs (CD26^+^)(Fig. 3G). We have recently shown that upon viral infection, inflammatory cDC2s (Fc*ε*RI+ cDC2s) arise, with capacities shared with both cDC1s (XCR1+) and cDC2s (CD172a+) (22). Upon SARS-CoV-2 infection, all DC subtypes were strongly present in the lungs *of Krt18*-hACE2 mice (Fig.3H-K). Interestingly, inf-cDC2s were already the most pronounced DC subtype at 4 dpi. Infl-cDC2s have been shown to be dependent on type I interferon signalling (22), and indeed both at 4 and 7 dpi, interferon-response genes such as *Isg15*, *Oas*, and *Cxcl9* were elevated in the lung of *Krt18*-hACE2 Tg mice. Collectively, SARS-CoV-2 infection induced pronounced pulmonary immune alterations in *Krt18*-hACE2 mice, including delayed neutrophil influx, loss of resident macrophages, and interferon-associated inflammatory cDC2 responses, whereas such changes were not observed in *Cdh5*-hACE2 mice.

### SARS-CoV-2 neuro-invasion occurs in *Krt18*-hACE2 but not in *Cdh5*-hACE2 mice

Given that microvascular injury and thrombosis have been reported in distal organs, including the brain, and that endothelial cells form a critical interface within the cerebral vasculature, we next investigated whether endothelial-restricted hACE2 expression permits SARS-CoV-2 neuro-invasion. Western blot analysis confirmed hACE2 protein expression in the brains of both hACE2 transgenic lines (Fig.4A). To quantify infectious viral particles, brain homogenates were subjected to plaque assays on VeroE6/TMPRSS2 cells (Fig.4B). Tissue samples were weighed post-collection to allow normalization, and assay sensitivity was defined by the Poisson 95% limit of detection (LOD□□), calculated from the total undiluted-equivalent volume plated. In female *Krt18*-hACE2 mice, SARS-CoV-2 tropism shifted toward the brain by 7 dpi, whereas no viral load was detected in male mice at this time point. Neuro-invasion was further confirmed by confocal imaging of brain sections stained with an anti-nucleocapsid antibody, revealing widespread viral signal throughout the brain of female *Krt18*-hACE2 mice at 7 dpi (Fig.4C). In contrast, SARS-CoV-2 could not be detected in the brains of *Cdh5*-hACE2 mice by any of the applied readouts, indicating that endothelial-restricted hACE2 expression alone is insufficient to support viral neuro-invasion.

**Fig 4:**
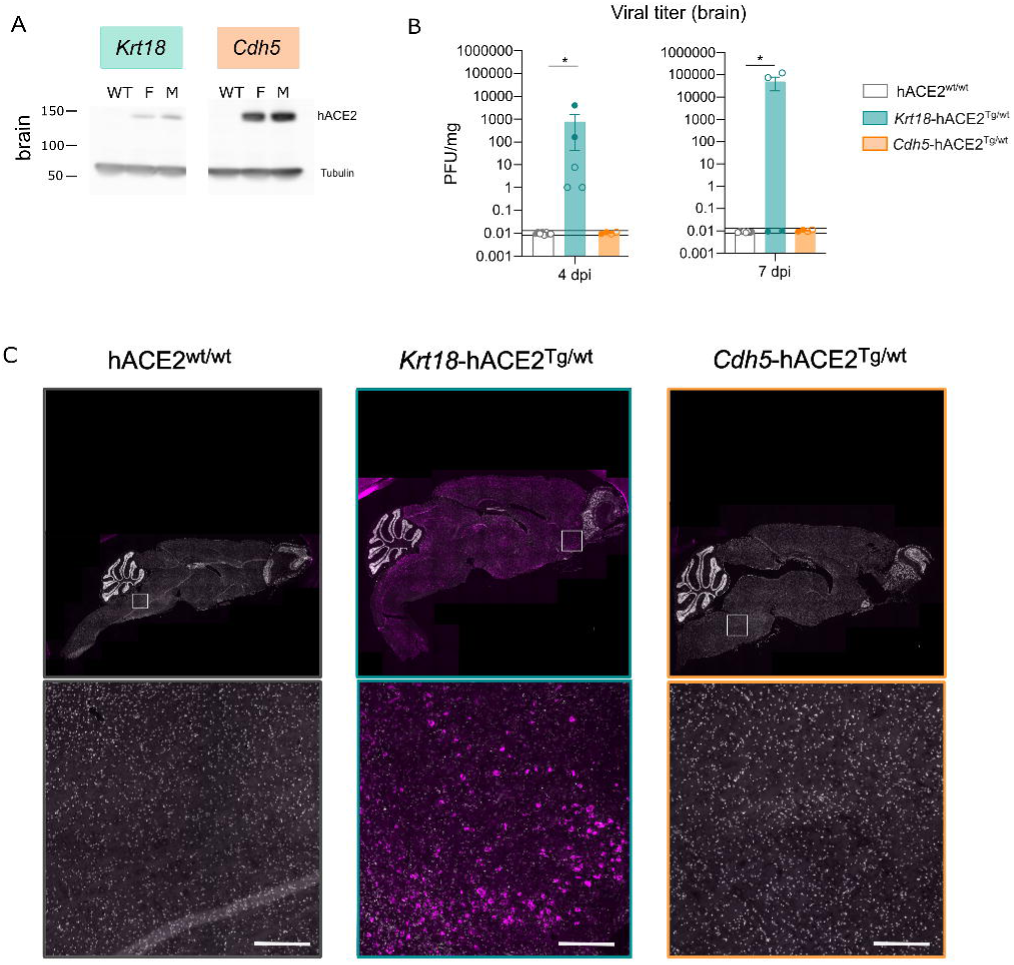
**A.** Western blot analysis of hACE2 protein in the brain of steady state hACE2-transgenic mice and accompanying β-tubulin loading controls. Samples were normalized to the same protein concentrations before loading on gel. **B:** Determination of infectious viral progeny by Plaque Assay on VeroE6/TMPRSS2 cells of homogenised brain samples of SARS-CoV-2 infected hACE2 Tg mice (i.t., 450 pfu) 4 and 7 days post infection (PFU/mg). PFU, plaque-forming units. Male mice are displayed in filled dots while females are represented with open dots. Black lines indicate the sample-specific plaque assay limit of detection converted to PFU/mg as LOD95%=4/M (M in mg; assay LOD = 8 PFU/mL; homogenization volume = mL). Nondetects are plotted at their own LOD. **C:** SARS-CoV-2 Nucleocapsid (N)-immunostaining (pink) in the brain of SARS-CoV-2 infected hACE2 Tg mice at 7 dpi. Scale bar = 200 *µm*. Statistics: After assessing normality by a Shapiro-Wilk normality test, non-parametric data were analysed with a Kruskal-Wallis test. Each transgenic line was compared to WT. ∗p < 0.05.

Collectively, these findings demonstrate that while SARS-CoV-2 robustly disseminates to and replicates within the brain of Krt18-hACE2 mice, endothelial-specific hACE2 expression does not confer susceptibility to neuro-infection, further supporting the notion that endothelial tropism alone is inadequate to drive SARS-CoV-2 pathology in distal organs.

## Discussion

From the earliest stages of the COVID-19 pandemic, it became evident that a substantial proportion of patients develop thromboembolic complications. Patients with severe COVID-19 display elevated levels of circulating endothelial cells and soluble markers of endothelial dysfunction, indicative of endothelial cell apoptosis and loss of vascular barrier integrity (23). Given that endothelial cells express ACE2, it was initially hypothesized that these vascular manifestations could result from direct viral infection of the endothelium. This concept was further popularized by electron microscopy studies reporting coronavirus-like particles within endothelial cells in tissues from COVID-19 patients (24). Despite these observations, convincing evidence for productive SARS-CoV-2 infection of endothelial cells *in vivo* remains limited. To date, neither human biopsy material nor animal models have definitively demonstrated efficient viral replication within endothelial cells (25–27).

This was the motivation to create *Cdh5*-hACE2 mice, with virus tropism restricted to endothelial cells. While pulmonary hACE2 expression and protein presence was confirmed, SARS-CoV-2 infection did not elicit clinical signs of infection, viral replication or histopathological changes. To bypass the requirement for epithelial barrier disruption and directly expose endothelial hACE2, Cdh5-hACE2 mice were also infected intravenously. Nonetheless, intravenous infection did not change the outcome. A limitation of the current study is that we did not comprehensively assess endothelial activation or dysfunction at the molecular level (VCAM1, ICAM1, and VE-cadherin expression), nor did we check pathways of coagulation, complement, or K-ras signalling.

These findings suggest that additional factors, most notably systemic inflammation, may be required to establish endothelial dysfunction in COVID-19. Endothelial cells are highly sensitive to inflammatory mediators generated during severe infection, and cytokine-driven endothelial activation is a well-recognized contributor to vascular injury. Pro-inflammatory cytokines such as IL-6, IL-1β, TNF-α, and MCP-1, together with complement activation, can amplify endothelial damage, promote thrombosis, and perpetuate multi-organ injury by reinforcing a vicious cycle of inflammation and vascular dysfunction (28–30).

An additional possibility is that SARS-CoV-2 may enter endothelial cells without establishing productive replication. In such a scenario, transient viral entry could initiate interferon and inflammatory signalling, while rapid degradation of viral RNA might render infection difficult to detect using conventional readouts.

An important consideration in interpreting endothelial involvement in SARS-CoV-2 pathology is the marked heterogeneity of endothelial cells across tissues. Recent single-cell transcriptomic studies have identified distinct endothelial phenotypes with immunomodulatory properties, including leukocyte recruitment, cytokine production, antigen presentation, and scavenger functions (31). Notably, lung endothelial cells are enriched for transcriptional programs associated with immunoregulation compared with endothelial cells from other organs (32). Within the pulmonary vasculature, a subset of capillary endothelial cells expresses high levels of genes involved in MHC class II–mediated antigen processing and presentation, suggesting a specialized role in immune surveillance against respiratory pathogens (31). These features highlight that pulmonary endothelial cells may be functionally distinct from endothelial populations in other vascular beds.

In parallel, endothelial susceptibility to SARS-CoV-2 is shaped by the expression of viral entry factors. Human endothelial cells generally express low levels of ACE2 and TMPRSS2, which likely limits productive infection (33). However, ACE2 expression varies across microvascular compartments and receptor availability can be dynamically modulated by inflammatory cues, indicating that endothelial permissiveness may be context-dependent (34). In addition, several alternative or auxiliary factors have been proposed to facilitate viral–endothelial interactions, including endosomal proteases such as cathepsins B and L, glycocalyx components such as heparan sulfate and sialylated glycoproteins, and receptors including neuropilin-1/2, vimentin, and CD147 (12). While these pathways may enhance viral binding or entry, their relative contribution to endothelial infection and pathology in vivo remains incompletely resolved.

In conclusion, our findings show that enforced endothelial-restricted SARS-CoV-2 tropism alone is insufficient to drive disease, challenging the notion that direct endothelial infection represents a universal primary pathogenic mechanism in COVID-19. At the same time, they leave open a potential role for endothelial infection in the context of multi-cellular viral spread and systemic inflammation. Future studies using these models may help elucidate how inflammatory environments modulate endothelial permissiveness to SARS-CoV-2 and contribute to vascular injury, thrombosis, and multi-organ failure in severe COVID-19.

## Material and Methods

### Mice

Mice were housed under specific pathogen-free conditions and used between 6 and 12 weeks of age. All experiments were approved by the animal ethics committee at Ghent University (EC2023-097) and were in accordance with Belgian animal protection law. *Krt18*-hACE2 transgenic mice were purchased from The Jackson Laboratory (strain ID 034860). *Cdh5*-hACE2 transgenic mice were generated in-house. A graphical representation of the construct used for the generation of this line can be found in Fig.1. In brief, the BAC clone RP23-277P19 containing the mouse *Cdh5* locus was ordered at BACPAC Resources Center. The FRT-kan/neo-FRT selection cassette was cloned downstream the pA sequence of pCAG-hACE2 using HindIII (partial digestion) and the resulting construct was used as a template for amplification of the recombination fragment. hACE2-pA-FNF was inserted into the start codon of *Cdh5* in exon 2 of the BAC via Red/ET recombination. The selection cassette was subsequently removed by expression of Flpe in bacteria. The modified BAC was digested with PacI and a 32 kB transgenic fragment was separated over PFGE, purified from gel and injected into fertilized oocytes. All founders were genotyped using primers specific to hACE2.

### SARS-CoV-2 infection

Mice were intratracheally infected with a sublethal dose of SARS-CoV-2 virus (450 pfu, SARS-CoV-2 D614G strain SARS-CoV-2/human/FRA/702/2020, obtained from the European Virus Archive (EVAG)). This virus was propagated on VeroE6/TMPRSS2 cells (NIBIOHN, JCRB1819) in Dulbecco’s Modified Eagle Medium (DMEM) supplemented with 2% fetal bovine serum (FBS). Two days after infection the growth medium was collected. After centrifugation for 10 minutes at 1000 g, the supernatant was collected, aliquoted and stored at –80°C. Virus titration was performed by plaque assay in which monolayers of VeroE6/TMPRSS2 cells were infected with dilutions series prepared in Dulbecco’s Modified Eagle Medium (DMEM) supplemented with 2% fetal bovine serum (FBS) in duplicate for 2 vials. Two hours after infection, Avicel was added to a final concentration of 0.3% (w/v). After 2 days of incubation at 37°C, the overlays were removed, and the cells were fixed with 3.7% paraformaldehyde (PFA) and stained with 0.5% crystal violet and counted. These experiments were performed in Biosafety Level 3 conditions at the VIB-UGent Center for Medical Biotechnology. All intratracheal treatments were given after mice were anesthetized with isoflurane (2 liters/min, 2 to 3%; Abbott Laboratories).

### Tissue sampling

Mice were euthanized by cervical dislocation or an overdose of pentobarbital (KELA Laboratoria) and organs were collected.

### Flow cytometry

Lung lobes were isolated, cut with scissors and then digested for 30 min in RPMI-1640 (Gibco, Thermo Fisher Scientific) containing 20 μg/ml liberase TM (Roche), 10 U/ml DNase I (Roche), and 5% of FCS (Bodinco) at 37°C. Next, lungs were filtered through a 70 μm cell strainer. Single cell suspensions were incubated with a mix of fluorescently labelled monoclonal antibodies (Ab) in the dark for 30 minutes at 4°C. To reduce non-specific binding, 2.4G2 Fc receptor Ab (Bioceros) was added. Dead cells were removed from analysis, using fixable viability dye eFluor506 or eFluor780 (Thermo Fisher Scientific). 123count eBeads Counting Beads (Thermo Fisher Scientific) were added to each sample to determine absolute cell numbers. Before acquisition, photomultiplier tube voltages were adjusted to minimize fluorescence spillover. Single-stain controls were prepared with UltraComp eBeads (Thermo Fisher Scientific) following the manufacturer’s instructions and were used to calculate a compensation matrix. Sample acquisition was performed on a NovoCyte Quanteon flow cytometer (Agilent) equipped with NovoExpress software. Graphical output was performed using FlowJo software (Tree Star, Inc.) and GraphPad Prism 10 (GraphPad Software, Inc.).

The following antibodies were used: anti-CD26 (FITC, BioLegend), anti-CD172a (PerCP-eFluor710, eBioscience), anti-CD317 (BV421, BD Biosciences), anti-CD11c (BV605, BioLegend), anti-CD88 (BV711, BD Biosciences), anti-CD19 (BV786, BD Biosciences), anti-CD40 (APC, BD Biosciences), anti-CD45 (AF700, BioLegend), anti-MHCII (APC-Cy7, eBioscience), anti-XCR1 (PE, BioLegend), anti-Fc*ε*RI (biotin, Life Technologies), streptavidin-PE-CF594 (BD Biosciences), anti-CD3 (PE-Cy5, BioLegend), anti-NK1.1 (PE-Cy5, BioLegend), anti-Ter119 (PE-Cy5, BioLegend), anti-CCR7 (PE-Cy7, BioLegend), anti-MHCII (FITC, eBioscience), anti-CD88 (PerCP-Cy5.5 BioLegend), anti-Ly6C (eFluor450, eBioscience), anti-CD11b (BV605, BioLegend), anti-Ly6G (BV650, BD Biosciences), anti-CD64 (BV711, BioLegend), anti-SiglecF (BV786, BD Biosciences), anti-CD45 (AF700, BioLegend), anti-bTCR (APC-eFluor780, BioLegend), anti-NK1.1 (PE, BD Biosciences), anti-CD11c (PE-TR, eBioscience), anti-CD4 (PE-Cy5, BioLegend), anti-CD8a (PE-Cy7, BioLegend). Spike-PE (see below).

### Recombinant S protein production

Recombinant SARS-CoV-2 S protein was produced and purified as previously described (58). The ectodomain of the SARS-CoV-2 S protein (ancestral Wuhan strain–derived, 2P, and mutated polybasic furin cleavage site) was expressed as a soluble, secreted protein, with a foldon-trimerization domain followed by a PreScission Protease cleave site, a His tag, and two StrepII tags (WSHPQFEK) fused to the C terminus. The protein was expressed by transiently transfected ExpiCHO cells and purified using a HisTrapHP column. For flow cytometry, S trimers were assembled into tetramers, yielding 12-nucleotide oligomer recombinant proteins. These were biotinylated and incubated with streptavidin (SAV)–conjugated phycoerythrin (PE) by the VIB protein core facility.

### Histology and confocal microscopy

During dissection, mice were transcardially perfused with PBS containing 0.002% heparin sodium followed by 1% PFA. After overnight fixation in 4% paraformaldehyde solution, lung and intestine were embedded in paraffin, cut into 5-μm slices, and stained with H&E. Images were taken with an AxioScan Slidescanner (Zeiss). For brain tissue, cryosections of 20 μm were cut and processed for immunostaining. For immunostainings, brain sections went through an extra round of fixation with 4% PFA, followed by blocking with 2% goat serum in PBS-T (PBS with 0.5% BSA and 0.3% Triton X-100) solution for 1 h at room temperature. The sections were then stained with primary antibodies (anti-Nucleocapsid-protein of SARS-CoV-2, SYSY, cat. HS-452 011) in blocking buffer at 4°C overnight. After washing with PBS, sections were stained with a AF633-conjugated goat-anti mouse secondary antibody (ThermoFisher Scientific) in blocking buffer for 2 h before washing and mounting. A Zeiss Axioscan Z.1 was used for imaging and images were processed using Zeiss Zen Software.

### RNA isolation, cDNA synthesis, and quantitative PCR (qPCR)

Lung samples were collected and mechanically disrupted using Qiagen TissueLyser beads and bead mill. RNA was isolated using TRIzol LS Reagent (Thermo Fisher Scientific) according to the manufacturer’s instructions. The cDNA was synthesized with the Sensifast cDNA synthesis kit (Bioline, Meridian Bioscience). qRT-PCR reactions were conducted in a LightCycler 480 (Roche) using the SensiFAST sybr no-ROX mix (Bioline). For the SARS-CoV-2 N-protein measurement Taqman PCR was performed using the LightCycler® TaqMan® Master (04535286001, Roche). Data were analyzed using qBASE software (BioGazelle).

Following primer pairs were used:

*hACE2* Forward:GGGATCAGAGATCGGAAGAAGAAA,

Reverse:AGGAGGTCTGAACATCATCAGTG.

*SC-2 N*

Forward: TTACAAACATTGGCCGCAAA

Reverse: GCGCGACATTCCGAAGAA

Probe: FAM-ACAATTTGCCCCCAGCGCTTCAG-BHQ1

*SC-2 S*

Forward: CCTTCCCAGGTAACAAACCAACC

Reverse: ACACACTGACTAGAGACTAGTGGC

*Ifn-γ*

Forward: ATGAACGCTACACACTGCATC

Reverse: CCATCCTTTTGCCAGTTCCTC

*Isg15*

Forward: GGTGTCCGTGACTAACTCCAT

Reverse: TGGAAAGGGTAAGACCGTCCT

*Oas1a*

Forward: GCCTGATCCCAGAATCTATGC

Reverse: GAGCAACTCTAGGGCGTACTG

*Cxcl9*

Forward: GTGGAGTTCGAGGAACCCTAG

Reverse: ATTGGGGCTTGGGGCAAAC

*Hprt*

Forward: TCCTCCTCAGACCGCTTT

Reverse: CCTGGTTCATCATCGCTAATC

*Rpl13a*

Forward: CCTGCTGCTCTCAAGGTTGTT

Reverse: TGGTTGTCACTGCCTGGTACTT

### Analysis of hACE2 protein expression by Western Blot

Organs were collected and mechanically homogenised using Qiagen TissueLyser beads and bead mill in RIPA buffer supplemented with cOmplete™ ULTRA protease inhibitor and PhosSTOP™ phosphatase inhibitor. The samples were vortexed and placed on a turner for 30 min at 4°C. Samples were centrifuged at 13000g for 10 min and the supernatant was used for Nano-orange protein measurement. Lysates were standardised to equal protein concentration and diluted in 4× Laemmli buffer containing 50□mM Tris-HCl pH 6.8, 2% sodium dodecyl sulphate (SDS), and 10% glycerol. Lysates were boiled for 10□min at 95□°C, proteins were separated by SDS-PAGE (8% gel) and transferred to a PVDF membrane. Immunoblotting was performed by overnight incubation at 4□°C using the anti-hACE2 primary antibody (ab108209, clone EPR4436, 1:1000, abcam) followed by a 2□h incubation at RT using the following secondary antibodies: HRP-linked anti-rabbit IgG (P0450, 1:1000, DAKO). As loading control, the blot was stained with anti-beta-tubulin-HRP (ab21058, abcam, 1:5000) 1,5 h on room temperature.

### Determination of viral titers by plaque assay

Lung and brain were directly collected in 2 mL CK14 and 7 mL CK28 Precellys ® tubes respectively. Tubes were weight before and after collection to determine the weight of the organ. Samples were homogenised in the Precellys® homogeniser for 2 x 10 sec at 5000 rpm, with a 15 sec break in between and spun down to remove debris. Monolayers of VeroE6/TMPRSS2 cells were infected with dilutions series of the samples prepared in FB medium:Opti-MEM™ in duplicate (FB medium: FluoroBrite DMEM medium (ThermoFisher Scientific cat# A1896702) supplemented with 5% FCS, 4 mM L-glutamine, NaPyruvate and P/S). One hours after infection, Avicel was added to the wells to a final concentration of 0.4% (w/v). After 2 days of incubation at 37°C, the overlays were removed, and the cells were fixed with 4% paraformaldehyde (PFA) and stained with 0.5% crystal violet.

## Statistical analysis

### Data information

Data are shown as means ± SEM. All data were analysed with a Shapiro-Wilk normality test to assess whether the data was normally distributed and whether a parametric or non-parametric test was applicable. For each Figure, the statistical tests are mentioned in the figure legends.

## Acknowledgements

We thank the VIB flow core and VIB transgenic core facility for their expert advice and service in this project. We thank Leen Vanhoutte and Tino Hochepied for their help in the generation of hACE2 transgenic mice. We thank Koen Sedeyn for providing training and service in the BSL3 facility at the VIB-UGent Center for Medical Biotechnology. We thank Prof. Roos Vandenbroucke for their analysis of histopathological brain sections. This publication was supported by the European Virus Archive GLOBAL (EVA-GLOBAL) project that has received funding from the European Union’s Horizon 2020 research and innovation programme under grant agreement No 871029.

## Funding

N.D. acknowledges support from an FWO Postdoctoral Fellowship - junior grant (3E003520).

B.S. acknowledges EOS joint programme of Fonds de la recherche scientifique—FNRS and Fonds wetenschappellijk onderzoek–Vlaanderen—FWO (G0H7518N EOS ID: 30981113 and G0H7322N EOS ID: 40007527).

S.D.C. is a predoctoral fellow supported by the Research Foundation-Flanders, Belgium (FWO-Vlaanderen 3S017320).

S.V. was a postdoctoral fellow of the Research Foundation-Flanders, Belgium (FWO-Vlaanderen; grant No. 3E002820) while working on this project and acknowledges support from FWO Vlaanderen senior research project grant No. 3G0A7422.

B.N.L. acknowledges support from FWO (3G0G4820) for this project and the Flanders Institute of Biotechnology (VIB).

